# NLRP10 Cleaves Oxidized DNA inhibited by OGG1 inhibitors: A Newly Identified Role in DNA Damage Processing and Senescence Regulation

**DOI:** 10.1101/2025.03.07.642110

**Authors:** Julia Elise Cabral, Sophia Lin, Haitian Zhou, Anna Wu, Angie Lackner, Minh Anh Pham, Feixuan Chi, Reginald McNulty

## Abstract

Mitochondrial DNA (mtDNA) release into the cytosol is a critical event in innate immune activation, often acting as a damage-associated molecular pattern (DAMP) that triggers inflammasome assembly. Here, we demonstrate that NLRP3 plays a direct role in cleaving and facilitating the release of D-loop mtDNA into the cytosol. We further show that NLRP3 interacts with NLRP10. NLRP10-mediated ox-DNA cleavage involves a Schiff base intermediate and is inhibited by small molecules known to inhibit glycosylases. These findings support a model where NLRP10 interaction with oxidized DNA may contribute to long-term senescence secretory phenotype and modulate inflammasome activation. Our study highlights a novel mechanism by which NLRP10 can respond to mitochondrial stress signals to influence innate immunity and suggests therapeutic potential for targeting these interactions in inflammatory diseases.

## INTRODUCTION

Mitochondrial DNA (mtDNA) serves as a potent damage-associated molecular pattern (DAMP) when released into the cytosol, triggering innate immune responses(1). Oxidized mtDNA (ox-mtDNA) accumulates under conditions of cellular stress, mitochondrial dysfunction, and aging, linking it to the activation of key immune pathways, including the cGAS-STING and inflammasome axes(2-4). Among these, the NLRP3 inflammasome plays a pivotal role in sensing mitochondrial distress and promoting IL-1β release(5, 6). Cryopyrin associated periodic syndrome (CAPS) gain-of-functions NLRP3 mutants have higher affinity for oxidized DNA(7, 8). However, the mechanisms governing mtDNA oxidation, processing, cytosolic release, and status of other NLR’s remain unclear and may be cell-type specific(9).

NLRP3 has been shown to co-IP with oxidized mitochondrial DNA(1) and cells lacking NLRP3 exhibit reduced IL-1β secretion upon oxidized mitochondrial DNA (ox-mtDNA) transfection(10). NLRP3 directly binds oxidized mitochondrial DNA with the pyrin domain preferentially binding ox-mtDNA(8). NLRP3 PYD has glycosylase-like activity enabling it to cleave oxidized DNA. Repurposed glycosylase inhibitors prevent both the interaction of NLRP3 with oxidized DNA and inflammasome activation(5).

NLRP3 is known to associate with mitochondria(11) and may regulate cleaved mtDNA D-loop release to cytosol via mPTP channels following inflammasome activation(2). This phenomenon implicates mtDNA in propagating inflammation and possibly reinforcing senescence-associated secretory phenotypes (SASP)(12). While inflammasome activation is classically associated with caspase-1-mediated cytokine maturation, emerging evidence suggests that the persistence of oxidized mtDNA in the cytosol may also contribute to prolonged inflammatory signaling and cellular senescence(13).

NLRP10, an enigmatic member of the NOD-like receptor (NLR) family which lacks an LRR domain, has been implicated in immune modulation, but its precise function remains controversial. Initial reports indicate that NLRP10 primarily acts as a negative regulator of NLRP3-dependent inflammation rather than forming a classical inflammasome itself(12, 14). However, recent studies suggest that mitochondrial damage stimulates NLRP10 to interact with ASC, contributing to inflammasome formation and IL-1b secretion independent of NLRP3 in keratinocytes(15).

Herein, we present evidence that NLRP3 and NLRP10 can associate. Moreover, NLRP10 binds and cleaves oxidized DNA, suggesting an alternative role in DNA damage processing. Structural comparisons indicate that NLRP10 harbors a glycosylase-like fold, typically associated with 8-oxoguanine DNA repair enzymes(16). We hypothesize that NLRP10 functions as a novel oxidized DNA-processing enzyme that regulates inflammatory signaling and senescence by modulating cytosolic DNA levels.

## RESULTS

### D-loop mitochondrial DNA release to cytosol is NLRP3-dependent

Drugs binding the NLRP3 pyrin domain have been shown to prevent NLRP3 from cleaving ox-mtDNA and inhibit both NLRP3 inflammasome activation and cytosolic mtDNA release(5). This suggests NLRP3 might directly play a role in ox-mtDNA release. We performed PCR using primers for cleaved D-loop mtDNA in mitochondrial and cytosolic fractions of immortalized macrophages(17, 18). PCR enabled detection of mtDNA large fragment, 5698bp, in the mitochondrial fraction for both wildtype and NLRP3 KO macrophages. Amplification of cleaved D-loop small fragment, 591bp, was found in the mitochondria of both wildtype and NLRP3 KO iBMDM’s. However, cytosolic D-loop mtDNA was found in activated wildtype macrophages, but completely absent in activated NLRP3 KO macrophages (**Fig. 1A**). The lack of cleaved D-loop mtDNA in NLRP3 deficient macrophages suggests NLRP3 directly interacts with the mitochondria to facilitate mtDNA cleavage and release.

**Figure 1:**
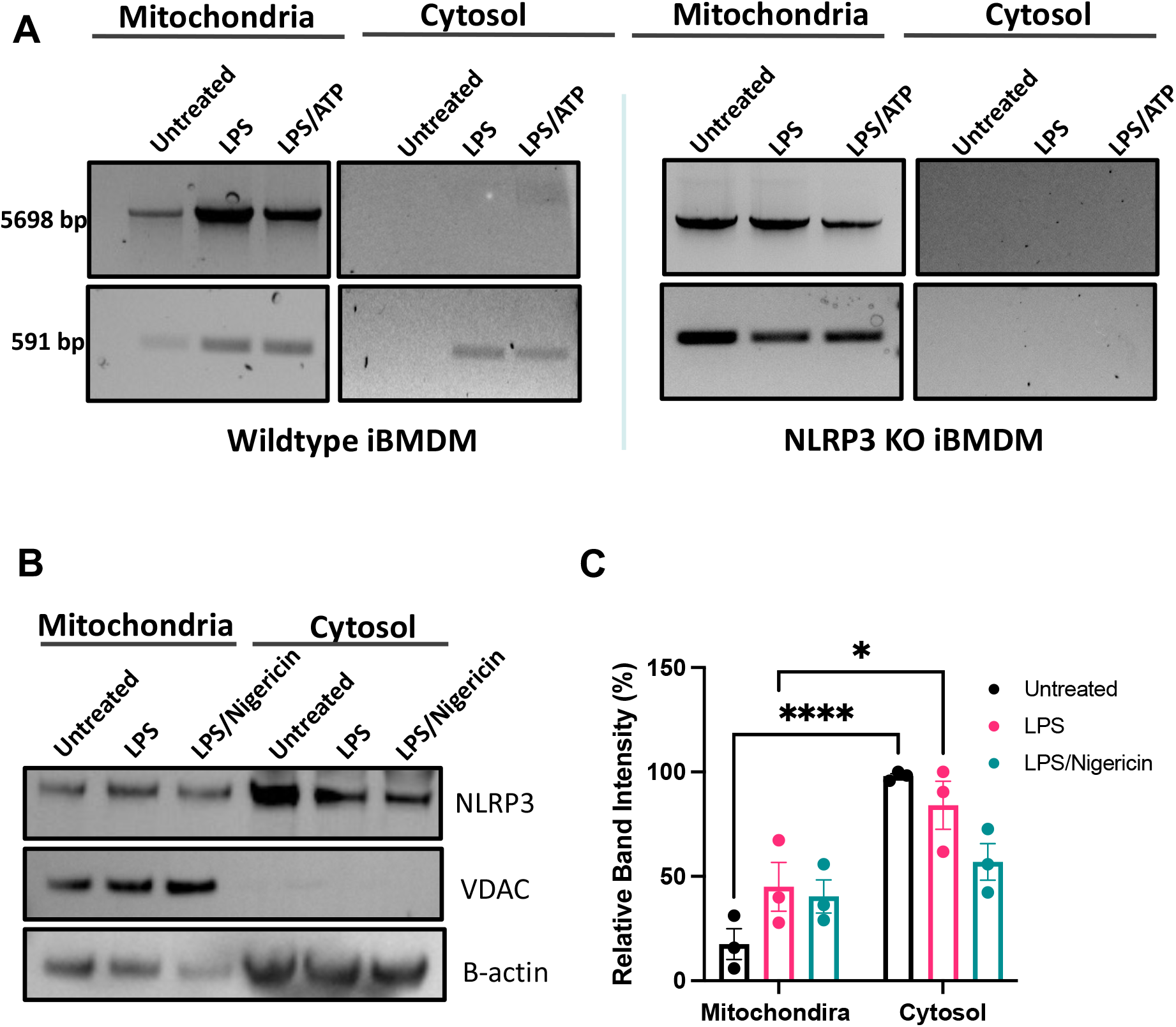
Mitochondrial DNA release is dependent on NLRP3. **A)** Mitochondrial and cytosolic fractions isolated from WT or NLRP3 KO iBMDMs treated with or without ATP. Amplification of 5698bp and 591bp fragments in both fractions are run on agarose gels. **B)** Western blot analysis of mitochondrial and cytosolic fractions isolated from THP-1 cells, probed for NLRP3 on a PVDF membrane. β-actin serves as a loading control for the cytosolic fraction, while VDAC confirms the absence of mitochondrial contamination in the cytosol. **C)** Quantification of NLRP3 in the mitochondria and cytosol with or without treatment with LPS and nigericin. Error bars: mean ± SEM, analyzed with two-way ANOVA. N = 3, *p=0.0256 **** p<0.0001.

### NLRP3 associates with mitochondria via MTS-like N-terminal helix

NLRP3 is activated by cardiolipin, a lipid found in the inner mitochondrial membrane. Upon stimulation, cardiolipin flips towards the outer membrane, potentially serving as a nucleation site to initiate NLRP3 inflammasome assembly. Since there are oftentimes variability in mouse compared to human cell immunology(19), we checked if NLRP3 could associate with the mitochondria in human monocytes. We purified the mitochondrial and cytosolic fractions from monocytes stimulated with LPS and LPS/nigericin. To be sure our isolations were contamination-free, we checked for the presence of voltage-gated anion channel, VDAC, in the cytosolic and mitochondrial fractions. VDAC, which is localized specifically in mitochondrial membrane, was only detected in the mitochondrial fraction and completely absent in the cytosol (**Fig. 1B-C**). We detected NLRP3 in both the mitochondrial and cytosolic fractions stimulated with either LPS or LPS/Nigericin. The presence of NLRP3 in the mitochondrial fraction is consistent with previous findings(11).

Although the fold of the death domain is shared amongst NLRP’s, they differ not only in percent identity, but the degree of structure/disorder at the N-terminus. Several software predictors of intrinsically disordered regions (IDR’s) identify the N-terminus of NLRP3 (and all other NLRP’s) as an IDR. This region could be the most subject to conversions between order and disorder(8). The N-terminal helix of NLRP3 PYD is required for both NLRP3 inflammasome activation and association with the mitochondria(11). We investigated if this helix fits canonical descriptions of mitochondrial targeting signals (MTS), which are often characterized by an amphipathic N-terminal helix(20). Using the program ChimeraX(21), we found that NLRP3’s N-terminal helix has one side that is hydrophilic with polar and charged amino acids while the opposite side was hydrophobic (non-polar). (**Fig. 2**). The inactive NLRP3 PYD N-terminal helix spanned amino acids 3-17. The N-terminus through the first helix_1-17_ was not predicted by DeepLoc(22) to associate with the mitochondria, with a mitochondria association probability of 0.5 (**Fig. 2A**). The N-terminus_1-16_ of the AlphaFold2 model of NLRP3 bound to oxidized DNA predicted a slightly higher mitochondrial association probability at 0.7 (**Fig. 2B**). However, analysis of NLRP3 bound to ox-mtDNA using SWISS Model predicted an N-terminal helix_1-11_ had mitochondrial probability localization score that was much higher at 0.9 (**Fig. 2C**). This analysis is significant because it supports NLRP3 may make minor changes to support interaction with the mitochondria membrane and oxidized mtDNA.

**Figure 2:**
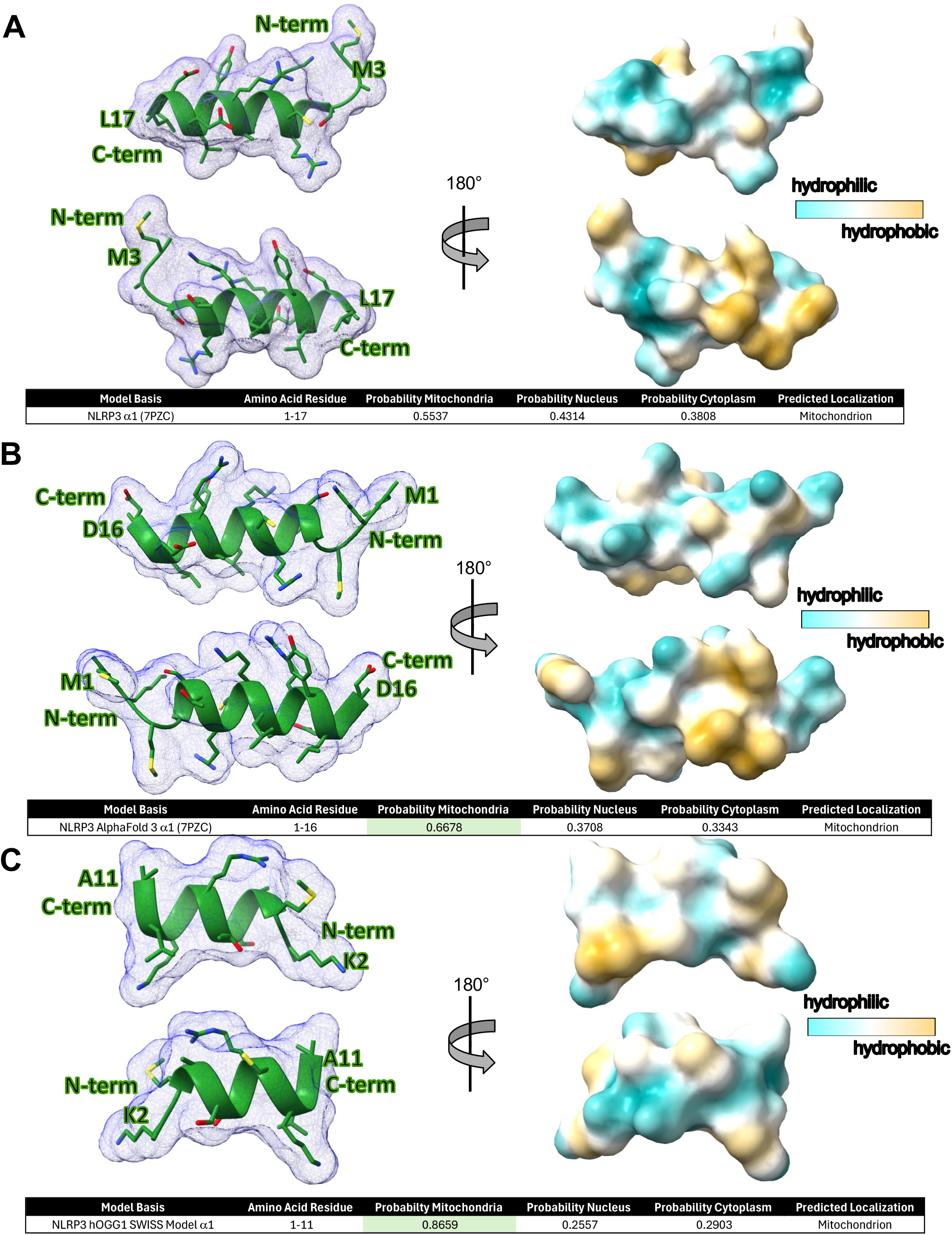
N-terminal NLRP3 helix is predicted to associate with the mitochondria. **A)** NLRP3 helix one_3-17_ is amphipathic with hydrophobic areas in orange and hydrophilic areas in blue (PDB 7PZC). **B)** NLRP3 pyrin domain AlphaFold3 model shows an amphipathic first helix_1-16_ with a predicted DeepLoc score of 0.6678. **C)** NLRP3 SWISS Model based on hOGG1 bound to oxidized DNA helix one_2-11_ shows regions of hydrophobicity and hydrophilicity. DeepLoc prediction software of N-terminal residues 1-11 predicts association with mitochondria with a score of 0.8659.

### NLRP3 associates with NLRP10

During NLRP3 activation ox-mtDNA exits the mitochondria and activates the NLRP3 inflammasome leading to an ASC speck in the cytosol. The presence of other pyrin domain containing proteins could be antagonistic if they are present at the same time. This has been seen for pyrin-only protein 1 (POP1), which inhibits NLRP3 inflammasome activation(23). Additionally, NLRP3 has been shown to co-localize with NLRP11 at ASC specks, where interaction between the two licenses inflammasome activation(24). We therefore checked if NLRP10 could interact with NLRP3. Using protein-G coated magnetic beads and a polyclonal antibody against NLRP3, we found NLRP3 could bind NLRP10 in THP-1 human monocytes (**Fig. 3A-B**). These results are significant because they suggest that the interaction between NLPR3 and ox-mtDNA may be modulated by the presence of NLRP10.

**Figure 3:**
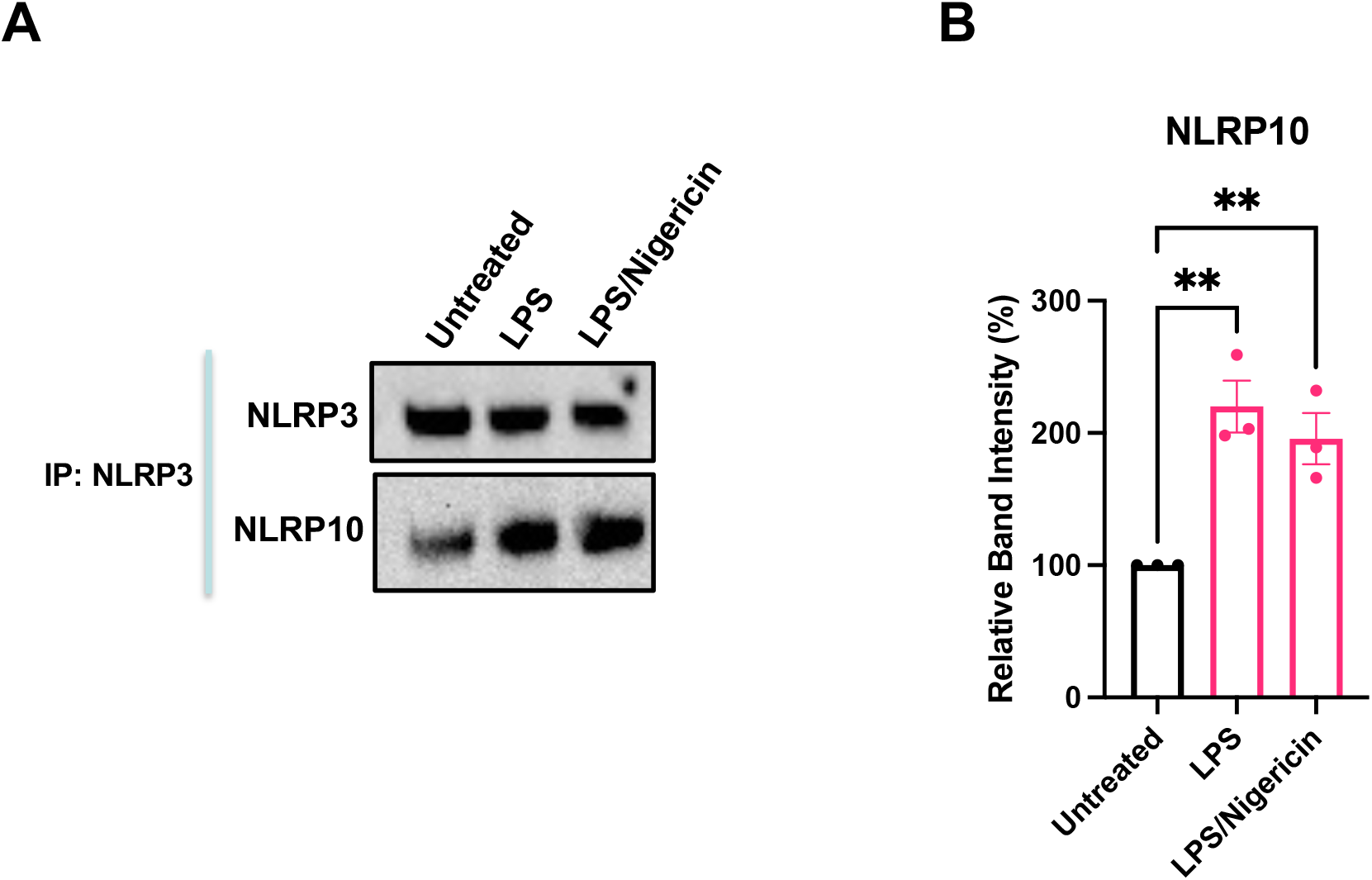
NLRP10 associates with NLRP3. **A)** Co-immunoprecipitation of NLRP3 pulls down NLRP10. Representative blots are shown. N=3. **B)** Quantification of NLRP10 band intensity. Error bars: mean ± SEM, analyzed with one-way ANOVA. N=3 **p = 0.0032 and 0.0095.

### NLRP10 has a glycosylase-like fold

NLRP3 is able to bind both non-oxidized DNA and ox-DNA similar to OGG1. Given the similarity of protein fold for NLRP3 and NLRP10 in the pyrin death domain fold, along with the identification of conserved Lys and Asp critical for nucleophilic attack, we wanted to address if NLRP10 could bind non-oxidized DNA. Recent structures of hOGG1 bound to non-oxidized DNA illustrate an interrogation complex where non-oxidized guanine is flipped into the active site, but uncleaved. Superposition of NLRP10 (1-102) with hOGG1 (248-325) bound to non-oxidized DNA(25) illustrated a similar protein fold (**Fig. 4A**). NLRP10 H1 and hOGG1 helix αL are roughly the same length at, 14.7 Å and 18.7 Å, respectively. These helices are not aligned. However, NLRP10’s H1 traverses the same direction and length of a disordered stretch of hOGG1. NLRP10 has a tight turn between Leu22 and Glu23. NLRP10 H2 traverses the same direction as OGG1. NLRP10 H2 has 4 turns with a length of 21.1 Å, while OGG1 αM has 3 turns and a length of 15.3 Å. Those helices are followed by a loop that turns the same direction. NLRP10’s H3 is a 1-turn helix in the middle of the loop. The fold matches up again at NLRP10 H4 and hOGG1 αN. hOGG1’s αN is 22.4 Å and has an extra turn, compared to NLRP10’s H4 which has a length of 19.2 Å. There is a tight turn for both proteins followed by similar lengths of NLRP10 H5 and OGG1 αO which both have lengths of 22 Å. Those helices are traversing the same direction and just out of phase relative to each other.

**Figure 4:**
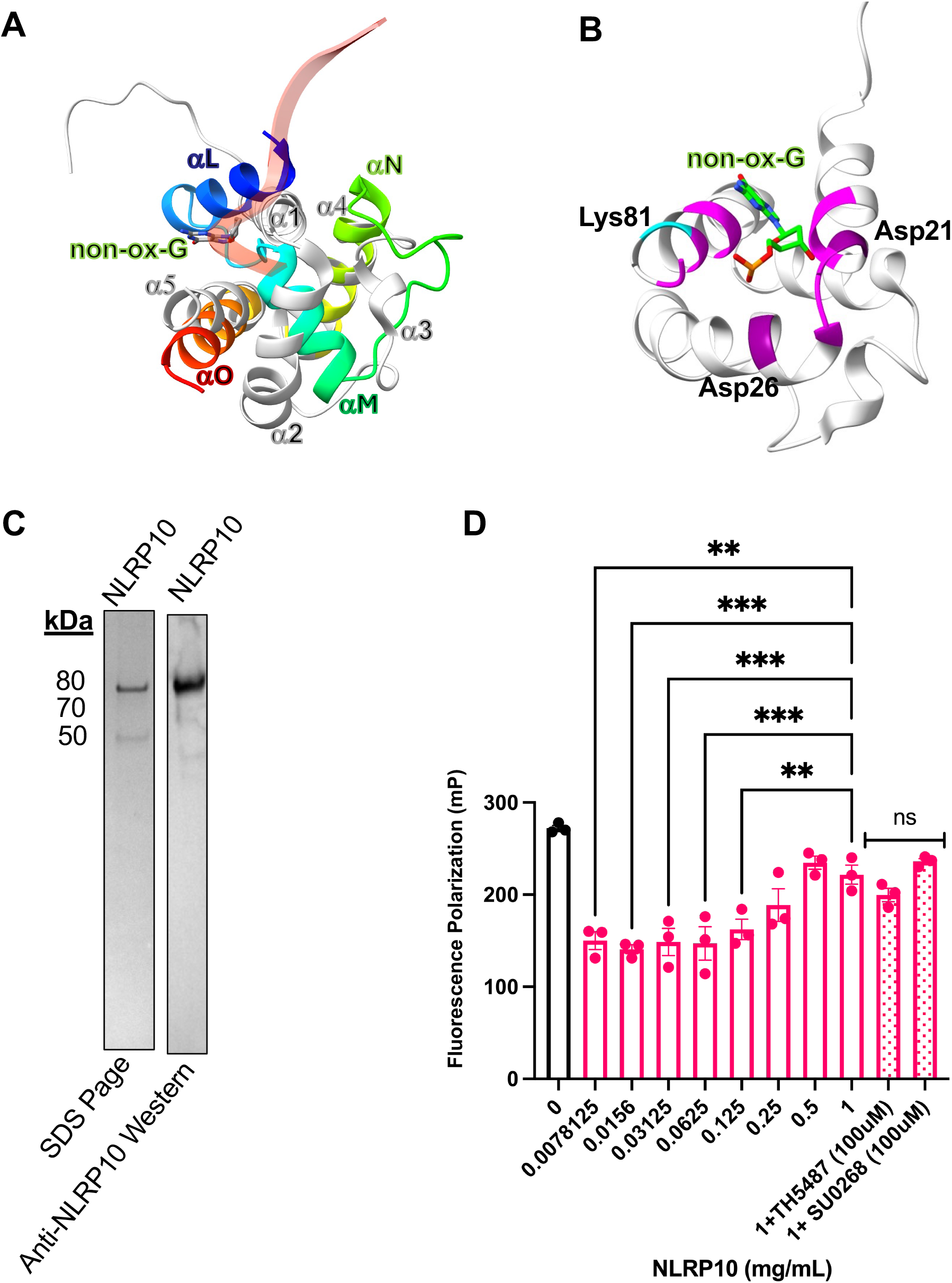
NLRP10 binds non-oxidized mitochondrial DNA. **A)** Superposition of NLRP10 (gray PDB 2m5v) with hOGG1 (rainbow PDB 8VX6) interrogating a non-oxidized guanine. **B)** Superposition from (A) with residues within 5 Å from non-oxidized guanine shown in magenta and lysine close to the catalytic site shown in cyan. **C)** SDS Page gel following flag column purification of NLRP10 (left). PVDF membrane probed for NLRP10 following FLAG column purification (right). **D)** Fluorescence polarization of cy5 labeled 20bp non oxidized DNA incubated with increasing concentrations of NLRP10. Error bars: mean ± SEM, analyzed with one-way ANOVA. N = 3, **p=0.0011, ***p=0.0003, 0.0009, 0.0007, **p=0.0072.

### NLRP10 binds DNA

A cryo-EM structure of OGG1 bound to nucleosome non-oxidized DNA illustrates a non-oxidized guanine flipped out from DNA and interacting with OGG1 active site(25). Superposition of NLRP10 with OGG1 bound to a flipped non-oxidized guanine illustrates a similar interrogation complex. The putative active site has Asp21, Asp26, and Lys81 that could potentially participate in nucleophilic attack and excise the guanine if it were oxidized (**Fig. 4B**).

Repurposed OGG1 inhibitors that prevent NLRP3 from binding oxidized DNA have no effect on NLRP3 binding non-oxidized DNA(25). So, we checked if NLRP10 could bind non-oxidized DNA and if the binding could be inhibited by repurposed inhibitors TH5487 and SU0268. We expressed and purified NLRP10 (75 kDa) via FLAG affinity tag (**Fig. 4C**). There was an initial decrease in FP signal upon adding 0.008 mg/ml NLRP10. However, the FP signal gradually increased with increasing concentrations of NLRP10. From 0.008 mg/ml to 0.5 mg/ml, the FP signal increased 36.1%, indicating NLRP10 could bind non-oxidized DNA (**Fig. 4D**). Addition of 100 mM TH5487 or SU0268 had no effect on binding non-oxidized DNA (**Fig. 4D**).

### NLRP10 binds and cleaves oxidized DNA

Given that NLRP10 could bind non-oxDNA and has a glycosylase-like fold (**Fig. 4A**), we examined if NLRP10 could interact with oxidized DNA and potentially cleave it. We first performed superposition of hOGG1 bound to oxidized guanine-containing DNA. The superposition had an RMSD of 0.895 Å (**Fig 5A**). The general path and alignment of the domain fold resembled that already shown in non-oxidized DNA. Residues potentially playing a role in catalysis within 5 Å of 8-oxo-dG included Asp26, Asp21, Lys78 and Lys 81(**Fig. 5B**).

**Figure 5:**
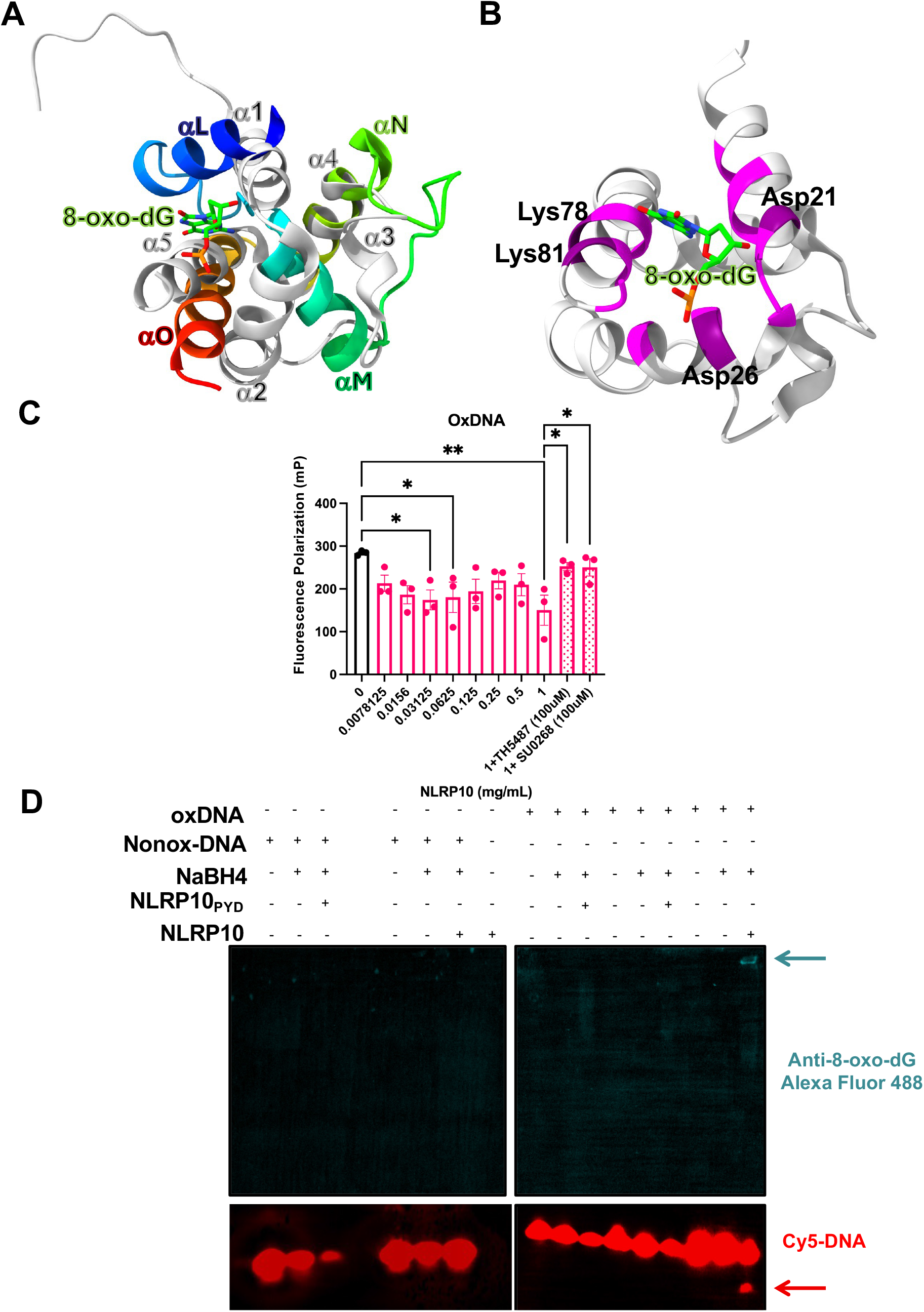
NLRP10 cleaves oxidized DNA. **A)** Superposition of NLRP10 (gray PDB 2M5V) with hOGG1 bound to 8-oxo-dG (rainbow PDB 1EBM). **B)** Superposition from (A) highlighting residues within 5 Å of oxidized guanine in magenta. Potential catalytic residues near the oxidized guanine are also indicated. **C)** Fluorescence polarization of cy5 labeled 20bp oxidized DNA incubated with increasing concentrations of NLRP10. Error bars: mean ± SEM, analyzed with one-way ANOVA. N = 3, *p = 0.0256, 0.0379, 0.0412, 0.0478, **p=0.005. **D)** Cy5 labeled 8-oxo-dG DNA incubated with either NLRP10_PYD_ or NLRP10 full length and sodium borohydride. Cy5 labeled DNA (red) anti-8-oxo-dG (teal). Cleaved oxDNA (red arrow). Covalent complex of NLRP10 bound to 8-oxo-dG (teal arrow).

Since NLRP3 has glycosylase-like activity, and the fold is similar between NLRP10 and OGG1, we examined if NLRP10 could cleave oxidized DNA. We incubated Cy-5 labelled oxidized DNA with increasing concentrations of NLRP10 to probe for enzymatic activity (**Fig. 5C**). The FP signal from DNA alone to 1 mg/ml NLRP10, decreased 47.1% percent, suggesting the DNA was being cleaved. FP measurements in the presence of TH5487 and SU0268 increased back to the baseline value and increased 40.6%. This illustrates the repurposed drugs prevent NLRP10 from cleaving oxidized DNA.

OGG1 glycosylase activity utilizes a conserved lysine that can perform a nucleophilic attack on 8-oxoguanine (26), which will nick DNA if it is double stranded, but cut the DNA if it is single stranded. Prior to cutting, the conserved lysine makes a Schiff base intermediate with the oxidized DNA. Structures of hOGG1 bound to oxDNA have been trapped by mutating K2Q or chemically by using sodium borohydride (NaBH_4_)(26). We incubated either non-oxidized or oxidized DNA with full length NLRP10 or only the pyrin domain of NLRP10 (NLRP10_PYD_) and checked for Schiff base formation and DNA cleavage. Non-oxidized DNA did not show any cleavage for the pyrin domain alone or full length (**Fig. 5D**). Cleavage was obtained with full-length NLRP10, as evidenced by an extra band underneath the primary oxDNA band in the lane (**Fig. 5D**). We investigated the mechanism of cleavage and found incubation of NLRP10 with sodium borohydride and ox-DNA led to a Schiff base shifted high in the gel when probed with an antibody against 8-oxo-dG Alexa Fluor 488. This result was significant because we observed both Schiff base formation and oxidized DNA cleavage in the same lane (**Fig. 5D**).

## DISCUSSION

We demonstrate that the release of mtDNA into the cytosol is dependent on NLRP3. The N-terminal helix of NLRP3 contains a predicted mitochondrial targeting sequence (MTS) and can interact with the mitochondrial membrane. To date, NLRP3 has not been found inside of mitochondria. But given the fact that the inner mitochondrial membrane inverts toward the outside and NLRP3 has an amphipathic-like N-terminal helix, it’s possible NLRP3 might be directly responsible for the flipping of the inner membrane. NLRP3 also interacts with NLRP10, and both proteins share a death domain with a structural fold akin to human glycosylase OGG1, exhibiting enzymatic activity specifically with oxidized, but not non-oxidized, mtDNA. To our knowledge, this is the first evidence illustrating that NLRP10 has glycosylase-like activity and can cleave oxidized DNA. Regarding the mechanism, boiling the samples and treating them with sodium borohydride (NaBH_4_) facilitated the formation of a Schiff base, which was trapped in a subset of samples. Meanwhile, a separate portion of the samples allowed simultaneous observation of NLRP10-mediated cleavage of oxidized DNA (**Fig. 5D**). These findings suggest a complex DNA-binding mechanism wherein non-oxidized DNA is differentially recognized compared to oxidized DNA. This conclusion is further supported by the observation that OGG1 inhibitors do not affect non-oxidized DNA but effectively inhibit enzymatic activity with oxidized mtDNA. Our data align with previous observations of NLRP3’s selective interaction with oxidized versus non-oxidized mtDNA(26).

NLRP10 activation has been linked to mitochondrial organelle damage(15). Interestingly, oblation of mtDNA still triggers NLRP10 activation. The mitochondrial inner membrane cardiolipin is likely to participate in the NLRP10 activation just like it does for NLRP3(27). So, it is likely either ligand is sufficient for NLRP10 activation.

Based on our findings that NLRP10 can cleave oxidized DNA similarly to hOGG1, it is plausible that NLRP10 functions as a sensor of oxidative damage, particularly in mitochondrial dysfunction. Given that mitochondrial oxidative stress and persistent DNA damage are hallmarks of cellular senescence(12), NLRP10 may play an underappreciated role in linking inflammasome activation to senescence-associated inflammation. It is possible that NLRP10 may detect oxidized DNA as a damage-associated molecular pattern (DAMP) and trigger a response that could promote senescence. The DNA itself could facilitate NLRP10 inflammasome activation, even though it is not required for activation. This is particularly relevant considering that inflammasomes, including caspase-1-dependent signaling, have been implicated in the senescence-associated secretory phenotype (SASP)(28), reinforcing the idea that NLRP10 could modulate the inflammatory microenvironment of senescent cells. Future studies should explore whether NLRP10 activation by oxidized DNA leads to persistent pro-inflammatory signaling and mitochondrial dysfunction, ultimately contributing to cellular aging and age-related pathologies. Or if NLRP10 cleavage of ox-mtDNA has a purpose to ameliorate the immune response.

Cellular senescence is a state of stable and irreversible cell cycle arrest that occurs in response to various stressors, including telomere shortening(29), oxidative stress(30), DNA damage(31), and oncogenic signaling(32). Senescent cells remain metabolically active but exhibit altered gene expression(33), resistance to apoptosis(34), and secretion of pro-inflammatory cytokines, growth factors, and matrix-remodeling enzymes known as the senescence-associated secretory phenotype (SASP)(12). While senescence initially serves as a tumor-suppressive and damage-control mechanism, the accumulation of senescent cells over time contributes to chronic inflammation, tissue dysfunction, and aging-related diseases.

Both IL-1β and ox-mtDNA are considered SASP factors. In the context of senescence, it depends on which is more predominant which would have more to do with tissue type and cellular environment. For example, in macrophages there is a high IL-1β dominant SASP, and ox-mtDNA-STING induced SASP is low(35). In epithelial cells, IL-1β SASP is low while ox-mtDNA/ cGAS/STING SASP is high(36). Dermal keratinocytes have high IL-1β SASP in response to UV, while also having high ox-mtDNA cGAS/STING SASP in response to photoaging(37).

Oxidized DNA, particularly from damaged mitochondria or nuclear sources, acts as a danger-associated molecular pattern (DAMP) that can be sensed by innate immune receptors, including NLRP3 and potentially NLRP10. When oDNA accumulates in the cytosol, it can activate inflammatory pathways such as the cGAS-STING axis and inflammasomes, leading to the production of type I interferons (IFNs), IL-1β, and IL-18, which are key components of SASP. This sustained inflammatory response reinforces senescence by inducing paracrine signaling, where neighboring cells are driven into a senescent-like state, thereby amplifying tissue dysfunction. Additionally, oxidative stress-induced DNA damage leads to persistent activation of the DNA damage response (DDR), maintaining the senescent phenotype and preventing cell proliferation. The failure to clear oxidized DNA or senescent cells can create a chronic inflammatory microenvironment, accelerating aging and promoting age-related diseases, including fibrosis, neurodegeneration, and cancer. This concept of inflammaging underscores how chronic, low-grade inflammation serves as a driving force in the progression of age-related pathologies(38). Given the connection between inflammasome activation initiated by pathogens(39), diet, and environmental toxicants future work will involve elucidating the molecular determinants of recognition for oxidized DNA and repurposed small molecules using cryo-EM(40-42).

## RESOURCE AVAILABILITY

We will be willing to distribute materials and protocols to qualified researchers in a timely manner.

### Lead contact

Further information and requests for resources and reagents should be directed to and will be fulfilled by the lead contact, Reginald McNulty.

### Materials availability

There were no unique materials generated by this study. Sources for all materials used to generate the data in this manuscript have been noted in the methods.

### Data and code availability

All data are available in the main text or the supplementary materials.

## ACKNOWLEDGMENTS

This work was supported by the National Institutes of Health NIAID grant T32AI177324 to J.E.C and K22AI139444 to R.M. We thank Michael Karin (UCSD School of Medicine) for providing immortalized macrophages.

## AUTHOR CONTRIBUTIONS

Conceptualization, R.M; methodology, J.E.C., S.L, H.Z, A.W, M.A.P, A.L.; Investigation, J.E.C, A.L, R.M.; writing—original draft, R.M. and J.E.C.; writing—review & editing, R.M. and J.E.C.; funding acquisition, R.M.; supervision, R.M.

## DECLARATION OF INTERESTS

The University of California Irvine is in the process of filling a patent with J.E.C., A.L., R.M. listed as inventors.

## METHOD DETAILS

### Immortalized bone marrow derived macrophage cell culture

Immortalized wild-type and knockout immortalized mouse macrophages(17) were provided by Micheal Karin at the University of California San Diego. Cell vials were thawed in a water bath at 37 °C and resuspended in prewarmed media (DMEM (Thermo Fisher) supplemented with 10% heat-inactivated FBS (sigma) and 1% penicillin-streptomycin (Thermo Fisher)). Cells were spun at 300 x g for 5 minutes at 4 °C to pellet the cells and the media and cryoprotectant were aspirated. The cells were then resuspended 5 mL of media and the cell viability and concentration were determined by diluting 10 μL of cells with 10 μL of Trypan Blue Stain (Thermo Fisher). The dilution was then added to a Countess Cell Counting Chamber Slide (Thermo Fisher) and evaluated using a Countess 3 FL Automated Cell Counter (Thermo Fisher). Cells were then further diluted to a final concentration of 0.25 × 10^6^ cells/mL and plated onto 175 cm^2^ culture flasks (Corning) where they were grown at 37 °C and 5% humidity. Cells typically doubled within 24–48 hours, after which they were scraped from the plate using Bio-One Cell Scrapers (Fisher Scientific), pelleted by centrifugation, counted, and expanded as previously explained.

### Immortalized bone marrow derived macrophage NLRP3 activation

Cells with a viability greater than 95% were diluted to a concentration of 0.75 × 10^6^ cells/mL, ensuring they would reach 1 × 10^6^ cells/mL by the following day. Cells were plated in 6-well plates (Corning) with 2 mL of cells per well. Cells were incubated overnight at 37 °C and 5% humidity. The next day, 4 μL (2 μL/mL) of 500 x LPS (Thermo Fisher) was added to each well and incubated for 4 hours. After incubation, LPS only wells were harvested, and 4 mM of ATP was added to each well for 1 hour. Cells were scrapped into 15 mL conicals and spun at 300 x g for 5 minutes. The supernatant was removed, and the cell pellet was washed with PBS and cell viability was checked. Cells were then spun at 500 x g and PBS was aspirated.

### Mitochondrial and cytosolic isolation in wild type/NLRP3 Knockout iBMDM and THP-1s

Protocol is adapted from a previous study(2). Briefly, following cell treatments, cell pellets were resuspended in 500 μL of cold mitochondrial extraction buffer (220 mM Mannitol, 70 mM sucrose, 20 mM HEPES-KOH, 1 mM EDTA, 2 mg/mL BSA, protease and phosphatase inhibitor cocktail tablet. Cells were passed through a 25-G syringe (BD Biosciences) 20 times on ice. The homogenized cell solution was then centrifuged at 1,000 x g for 15 minutes at 4 °C to pellet cell debris. The supernatant was then centrifuged at 12,000 x g for 15 minutes to pellet the mitochondria. The supernatant was removed and labeled as the cytosolic fraction. The mitochondrial pellet was resuspended in cold 300 μL of cold RIPA buffer. Bradfords were performed on both fractions with the Quick Start Bradford Protein Assay Kit (BioRad) prior to western blotting. NuPAGE 4-12% gels (Thermo) were run, and gels were transferred onto PVDF membranes. Membranes were blocked in 2.5% BSA and probed with pyrin targeting NLRP3 (Adipogen–AG-20B-0014-C100), VDAC (Cell Signaling - #4661), and β-actin (Santa Cruz - #47778).

### Amplification of 591 base pair and 5698 base pair fragments from cytosol and mitochondria of mouse iBMDMs

The mitochondrial and cytosolic fractions from immortalized bone marrow derived macrophages were isolated as described above. The mitochondrial pellet was lysed using the lysis buffer from the Quick-DNA/RNA Miniprep Kit (Zymo Research), while the cytosolic fraction was used without further processing. DNA was then purified with the same kit, and the concentrations were measured using a NanoDrop spectrophotometer. PCR was performed to amplify both the large (5698 bp) and small (591 bp) fragment as previously described(2). Briefly, PCR reactions were set up using the Phusion Hot Start 2x Master Mix (NEB #M0536) per the manufacturer’s protocol. For 5698 bp PCRs on the cytosolic fractions, 50 ng of DNA was used as the template. For all other reactions, 5 ng was used as the template. The following PCR conditions were used: hot start at 98 °C for 3 minutes, melting temperature of 98 °C for 10 seconds, annealing temperature of 60 °C, and extension temperature of 72 °C for 30 seconds (591 bp) or 3 minutes (5698 bp), followed by 72 °C for 20 minutes and kept at 4 °C. The PCR products were then analyzed on 1% agarose gels stained with ethidium bromide.

### Fluorescence polarization

A 100 μM stock of Cy5-labeled 20 base pair DNA (oxidized or non-oxidized (IDT)) was diluted 1:100 in DEPC-treated water. Protein was serially diluted eight times with binding buffer (50 mM Tris HCl (pH 7.4), 100 mM NaCl, 2 mM MgCl_2_, 12% glycerol) starting from the highest concentration of 1 mg/mL to the lowest concentration of 0.0078125 mg/mL. ATPγS was added to the reaction at a final concentration of 1 mM. A 10 μL aliquot of 1 μM DNA stock was added to each tube and incubated on ice in the dark for 1 hour. For samples containing TH5487 and SU0268, the protein was pre-incubated with protein for 1 hour on ice prior to DNA addition. 10 μL of stock DNA, 80 μL of buffer, and 1 mM ATPγS was added as a control. After incubation, the incubation was divided into triplicate and 30 μL of reaction volume was added to three wells of a black-walled, clear-bottomed 384-well plate (Agilent #204623). Fluorescence polarization was then read using a BioTek Synergy H1 plate reader with Generation 6 software (Agilent) at an excitation of 620/40 and an emission of 680/30, specific to Cy5. The polarization values were calculated based on the parallel and perpendicular light and G-factor-corrected.

### NLRP10 pyrin and full-length protein expression and purification

NLRP10 plasmid was a gift from Thomas Kufer (Addgene plasmid # 38141). For the NLRP10 pyrin domain, site directed mutagenesis was performed (NEB #E0554S) to introduce a stop codon after the 100^th^ amino acid. The plasmids were sequenced, grown at large scale in DH5α cells, and then the DNA was purified with PureLink HiPure Plasmid Maxiprep kit (Thermo Fisher). The proteins were each expressed in 500 mL of media using the Expi293 Expression System (Thermo Fisher). Briefly, cells were grown to roughly 3×10^6^ cells per milliliter and ≥95% live cells. Enhancers were added 16 hours after transfection. Once the cells reached a viability of <80% live cells, they were harvested by spinning at 1200 rpm in a swinging bucket rotor (JS-4.750, Beckman). After centrifugation, the media and dead cells was aspirated, and the pellet was washed with cold PBS to remove residual media. The cells were then harvested again by centrifugation and the PBS was aspirated. The protein was purified on the AKTA Avant. Cells were resuspended in lysis buffer (1 mm PMSF, 50 mM Tris HCl pH 7.4, 300 mM NaCl, 0.1% SDS, 10% glycerol, 1% Triton X-100, protease inhibitor tablets (Sigma)). The lysate was then sonicated for 42 seconds in intervals of 2 seconds on and 8 seconds off. The lysate was then clarified by spinning at 100,000 x g for 1 hour and subsequently passed through a 0.45 μm filter prior to affinity chromatography. The chromatography column (XK16) was packed with flag resin and equilibrated with binding buffer (150 mM NaCl, 50 mM Tris, 0.05% NP-40). The protein was eluted off with elution buffer (0.1 M glycine, 0.08% NP-40) into a 96 well 2 mL fraction collector containing Tris HCl at a final concentration of 60 mM. Fractions were analyzed on a total protein NuPAGE 4–12% Bis-Tris gel run at 200 V for 30 minutes. Fractions were further analyzed with PVDF membrane western blots blocked in 2.5% BSA and probed with NLRP10 NACHT targeting antibody (Cell Signaling) at a dilution of 1:1,000. Based on the results from the gels and westerns, similar fractions were pooled and concentrated in a 50 molecular weight cut off spin concentrator (Sigma). The protein was then dialyzed in 500 μL 10 kd dialysis bags (Sigma) for 4 hours at 4 °C into dialysis buffer (20 mM Tris HCl, 200 mM NaCl, 0.08% NP-40, 10% glycerol, 1 mM DTT).

### ChimeraX and hydrophobicity maps

The inactive structure of NLRP3 (PDB 7PZC)(43) was opened in ChimeraX. The N-terminus through the end of the first helix was selected (3-17). A separate .pdb file was saved to isolate this helix and the hydrophobicity was displayed. To generate the AlphaFold model, the sequence of NLRP3’s pyrin domain (PDB 7PZC, amino acids 1-91) was inputted into AlphaFold3 using the Colab server. Next, the sequence of single-stranded oxDNA (PDB 1EBM, Chain C) was added to the same query. Alphafold generated five putative models, and the model with the highest combined confidence score was evaluated using ChimeraX. The N-terminus through the end of the first helix was selected (1-16) and the hydrophobicity displayed on the isolated helix. To generate the SWISS Model, the amino acid sequence of hOGG1_(249-325)_ that aligned with the NLRP3 pyrin_(1-91)_ domain was saved as a.pdb file. The sequence was uploaded to SWISS-Model in their User Template Modeling input option as the template file. Then, NLRP3 amino acids 1–85 were loaded as the target file (amino acids 1–90 did not generate a model). The SWISS-Model projection produced one model of NLRP3 based on the template hOGG1 structure. The N-terminus through the end of the first helix (2-11) was isolated in ChimeraX and the hydrophobicity shown.

### THP1 cell culture

THP-1 cells were purchased from Invivogen. Cells were thawed in a water bath and resuspended in 5 mL of RPMI 1640 without phenol red (Thermo Fisher), supplemented with 1× non-essential amino acids (Thermo Fisher), 1× Penicillin-Streptomycin-Glutamine (Thermo Fisher), 1× sodium pyruvate (Thermo Fisher), and 10% heat-inactivated FBS (Thermo Fisher). The cells were centrifuged at 300 × g for 5 minutes, and the cryoprotectant and media were aspirated. Cells were then resuspended in 5 mL of media, counted using a Countess 3 FL Automated Cell Counter (Thermo Fisher), and diluted to 0.25 × 106 cells/mL. Cells were incubated at 37 °C and 5% humidity

### THP1 NLRP3 activation assay

THP-1 cells with a viability greater than 95% were diluted to a concentration of 0.75 × 106 cells/mL the day before the assay was performed. Cells were plated in 6-well plates (Corning) with 2 mL of cells per well. Cells were incubated at 37 °C and 5% humidity. Once cells reached 1 × 106 cells/mL, 4 μL of LPS (Thermo Fisher) was added to each well and incubated for 16 hours. After incubation, LPS wells were harvested, and 10 μM nigericin was added to each well for 1 hour. Cells were transferred to 15 mL conical tubes, spun at 300 x g and the supernatant was saved for western blot analysis. Cells were washed with cold PBS and cell viability was determined as described above. Cells were then spun at 500 x g and PBS was aspirated.

### Co-immunoprecipitation

THP-1 cell pellets were lysed with gentle lysis buffer (50 mM Tris HCL, 150 mM NaCl, 5 mM EDTA, 1% NP-40, 1 mM PMSF). Bradfords were run on cell lysates (BioRad) to ensure equal loading onto the beads. PureProteome Protein G magnetic beads (Sigma) were resuspended and 50 μL were removed for each replicate. Tubes containing beads were placed on a magnet and the storage puffer was discarded. Beads were washed with 500 μL of wash buffer (PBS supplemented 0.1% tween). Beads were resuspended in wash buffer and incubated with polyclonal NLRP3 antibody (ABclonal A12694) for 10 minutes at room temperature. Beads were placed on the magnet and washed with wash buffer to remove unbound antibody. Equal amounts of protein were incubated with the beads overnight at 4 °C. Beads were pulled back and unbound sample was removed. Beads were washed with wash buffer 3 times. NuPAGE sample buffer and reducing agent (Thermo Fisher) were added at 1x and boiled for 10 minutes at 80 °C. Beads and sample were loaded together onto NuPAGE 4-12% gels (Thermo Fisher). Gels were transferred onto PVDF membranes, blocked with 2.5% BSA, and probed for NLRP3 (1:10,000 Adipogen AG-20B-0014-C100) and NLRP10 (1:1,000 Cell Signaling E5×5A).

### Schiff Base trapping of NLRP10 and oxidized DNA

The protocol was adapted from a previous study(44). Briefly, wild type NLRP10 or NLRP10_1-100_ was added to a reaction buffer containing 50 mM NaBH4, 25 mM potassium phosphate (pH 6.8), 1 mM DTT, 1 mM EDTA at their final concentrations. A 100 μM stock of Cy5-labeled 20 base pair DNA (oxidized or non-oxidized (IDT)) was diluted 1:100 in DEPC-treated water. 5 μL of DNA was added to each tube and incubated at 37 °C for 30 minutes, followed by placement on ice for 1 hour. Readouts were performed immediately after incubation or several days later following incubation at 4 °C. Samples were analyzed on 4-20% Tris-Glycine gels (Thermo Fisher). Samples were mixed with orange loading dye (NEB) containing SDS, boiled at 70 °C for 10 minutes, and loaded onto the gel. Fluorescent signals were detected both after electrophoresis and following transfer onto a PVDF membrane. Oxidized DNA signals were further analyzed via western blot after UV crosslinking of the membrane for 5 minutes. Blots were blocked with 2.5% BSA at room temperature and probed with an 8-oxo-dG antibody (1:1,000, Rockland). The secondary antibody, Alexa Fluor 488 goat anti-mouse IgG/IgM (H+L) (1:2,500, Thermo Fisher), was incubated overnight at 4 °C, and membranes were imaged using iBright software.

## QUANTIFICATION AND STATISTICAL ANALYSIS

A one-way or two-way ANOVA was used to conduct all statistical analyses herein. All statistical analyses were performed as indicated in the Figure legends where N represents the number of replicates. The data all represent mean ± SEM where p-values <0.05 were considered statistically significant.

